# Discovering SARS-CoV-2 neoepitopes and the associated TCR-pMHC recognition mechanisms by combining single-cell sequencing, deep learning, and molecular dynamics simulation techniques

**DOI:** 10.1101/2023.02.02.526761

**Authors:** Kaiyuan Song, Honglin Xu, Yi Shi, Jie Hao, Lin-Tai Da, Xin Zou

**Author notes:** Corresponding author (XZ); (LTD); (JH).

## Abstract

The molecular mechanisms underlying the recognition of epitopes by T cell receptors (TCRs) are critical for activating T cell immune responses and rationally designing TCR-based therapeutics. Single-cell sequencing techniques vastly boost the accumulation of TCR sequences, while the limitation of available TCR-pMHC structures hampers further investigations. In this study, we proposed a comprehensive strategy that incorporates structural information and single-cell sequencing data to investigate the epitope-recognition mechanisms of TCRs. By antigen specificity clustering, we mapped the epitope sequences between epitope-known and epitope-unknown TCRs from COVID-19 patients. One reported SARS-CoV-2 epitope, NQKLIANQF (S_919-927_), was identified for a TCR expressed by 614 T cells (TCR-614). Epitope screening also identified a potential cross-reactive epitope, KLKTLVATA (NSP3_1790-1798_), for a TCR expressed by 204 T cells (TCR-204). According to the molecular dynamics (MD) simulations, we revealed the detailed epitope-recognition mechanisms for both TCRs. The structural motifs responsible for epitope recognition revealed by the MD simulations are consistent with the sequential features recognized by the sequence-based clustering method. This strategy will facilitate the discovery and optimization of TCR-based therapeutics. In addition, the comprehensive strategy can also promote the development of cancer vaccines in virtue of the ability to discover neoepitopes and epitope-recognition mechanisms.

## Introduction

Triggered by the recognition of antigens derived from pathogens or tumor-associated mutations, the T cell immune response is integral to the adaptive immune system for immune surveillance and clearance[1-3]. T cell receptors (TCRs), as heterodimers on the surface of T cells, take charge of recognizing antigenic peptides presented by the major histocompatibility complex (MHC, also termed human leukocyte antigen or HLA in humans) on the surface of antigen-presenting cells to activate T cell responses[3]. During the development of T cells in the thymus, TCR genes are generated by V(D)J recombination, thereby different germline gene usages and imprecise gene segments joining endow the TCR sequences with enormous diversity[4, 5]. It is estimated that approximately 2×10^19^ αβTCR sequences can be generated in humans[6], although only a fraction of which are present in an individual[7, 8]. Accordingly, the TCR repertoire, owing to its natural diversity, bears the potential to recognize various antigenic peptides. In addition, the polyspecificity that a certain TCR is capable of recognizing multiple distinctive peptide-MHC (pMHC) ligands further broadens the antigenic peptide repertoires under immunosurveillance[9, 10]. On the other hand, the tremendous TCR sequences and polyspecificity complicate the mechanistic investigation and limit therapeutic applications.

Harnessing T cell immune responses by engineered TCRs or TCR-based molecules is a promising means of immunotherapy; however, the inadequate understanding of epitope recognition by TCRs impedes the comprehensive utilization of immunological weapons[11]. With next-generation sequencing and single-cell approaches exploited for elucidating T cell immune responses[8, 12], innumerable data on TCR sequences and T cell gene expression have been released. Consequently, multiple methods used for TCR repertoire analysis were developed to understand T cell immune responses and assist clinical applications[7, 13, 14]. The diversity originally used for quantifying the distribution of species in ecology has become a general measure to describe the TCR repertoire[14]. By delineating the features of TCR repertoires from individuals in various contexts, it is accessible to predict the immune status associated with diseases[15-17] and the responses to immunotherapy[18]. Despite various high-throughput sequencing techniques developed, it remains an arduous task to obtain epitope information for TCR repertoires[7, 14]. To infer the antigen specificity, two seminal studies have deployed the sequence similarity to cluster TCRs[19, 20]. According to sequence-based clustering methods, the TCRs falling into the same cluster share similar antigen specificities. Thereafter, several specificity clustering methods were developed and utilized for disease-associated TCR identification[21-25]. Meanwhile, TCR sequencing techniques with antigen specificity[26] and VDJdb[27], a curated database storing tens of thousands of epitope-known TCR sequences, provide valuable resources for clustering-based specificity analysis. These methods and resources, to some extent, promote the investigation of epitope recognition mechanisms at the sequence level.

Compared with the rapidly increasing data generated by high-throughput sequencing, the limited structural data of TCR hampers the investigation of TCR recognition mechanisms and further translational applications, such as the design of TCR-based therapeutics[11]. According to the structural T cell receptor database (STCRDab[28]), only ∼600 TCR-associated structures are available, which is far too less compared with the immense diversity of TCR repertoires[29]. To overcome the limitation of structural data, computational tools have been developed and applied to the design of TCRs[30-33]. Assisted by the development of deep learning-based methods in recent years, the protein structure in apo form can be readily obtained by AlphaFold or RoseTTAFold[34, 35]. To construct the complex structure, one can further employ information-driven molecular docking to predict the binding mode of protein to its ligands, e.g., for antigens and antigen receptors[36, 37]. Therefore, researchers could potentially obtain the TCR-pMHC ternary structure by combining sequence analyses and model constructions, thereby revealing the molecular mechanisms underlying the activation of T cell immune responses in silico[3, 38]. However, due to the lack of a comprehensive strategy integrating single-cell sequencing data with structural information, it remains a question whether the structural or kinetic properties of TCR-pMHC interactions are associated with the cellular characteristics of T cells captured by single-cell techniques.

To advance the investigation of epitope recognition by TCRs, it is necessary to develop a comprehensive strategy that leverages large sequence data and structure-modeling tools. In this study, we proposed a computational pipeline to identify disease-associated TCR-pMHC complexes and unveil the specific interacting partners responsible for epitope recognition. Combining epitope-unknown and epitope-known TCRs associated with SARS-CoV-2, we mapped the epitope information for epitope-unknown TCRs using GLIPH[20], a sequence-based TCR clustering software. We also exploited similarity searching and immunogenicity prediction to discover potential epitopes. A reported SARS-CoV-2 epitope from the spike protein, NQKLIANQF (S_919-927_), and a potential cross-reactive epitope from the nonstructural protein 3 (NSP3), KLKTLVATA (NSP3_1790-1798_), were identified for two TCRs expressed by highly expanded T cells. We further performed molecular dynamics (MD) simulations for the identified TCR-pMHC complexes and pinpointed the critical structural motifs in TCRs responsible for epitope recognition. Our computational strategy bridges the single-cell sequencing data of TCRs, epitope sequences, and the structural dynamics of TCR-pMHC, providing a means to obtain TCR-pMHC interactions at the atomic level. This strategy can facilitate future attempts to design TCR-based therapeutics and cancer vaccines.

## Methods

### Datasets collection

The epitope-unknown TCR data were collected from a massive single-cell dataset sampled from healthy controls and COVID-19 patients[39] (NCBI GEO database: GSE158055). The collected data mainly included germline gene usages, the nucleotide and amino acid sequences of the complementarity-determining regions 3 (CDR3s), and the cell type of originated cells. For convenience, we reannotated the cell subtypes according to marker gene expression (Table S1) and retained only αβTCRs, resulting in 213,755 epitope-unknown TCRs (Table S2). In addition, 43,252 epitope-known human TCRs or TCR β chains were collected from the VDJdb database[27].

### TCR diversity analysis

Samples that contained more than five distinctive TCRs that are different in germline gene usages or nucleotide sequences of CDR3s were retained for the diversity analysis. For each subtype of T cells, the TCR diversity was calculated as Shannon’s entropy[39]:

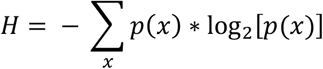

where p(x) represents the frequency of the TCR.

### Antigen specificity clustering

For the epitope-unknown TCRs, 8,507 unique TCRs that occurred more than once were extracted to make up the UNK dataset. The epitope-known TCR datasets VDJ-S and VDJ-N, derived from the VDJdb database, contained 1,766 TCRs targeting SARS-CoV-2 epitopes and 29,101 TCRs targeting antigens from other species, respectively. The TCRs from the UNK dataset were clustered with TCRs from the VDJ-S and VDJ-N datasets, respectively. The clustering processes were performed using the GLIPH (grouping lymphocyte interactions by paratope hotspots) algorithm[20]. According to the clustering algorithm, the CDR3β sequences of TCRs in the same cluster show either the global similarity that only one amino acid is different or the local similarity that enriched sequence motifs exist. However, due to the global similarity, the highly distinctive TCRs that do not share antigen specificities can be grouped in the same cluster via the connections between similar TCRs. To improve the clustering accuracy, we trimmed clusters by retaining the epitope-known TCRs that were different in only one position or showed local similarity with epitope-unknown TCRs, and vice versa. Finally, the epitope information was mapped between epitope-known and epitope-unknown TCRs in the same cluster.

### Analysis of TCR-pMHC crystal structures

A total of 133 human TCR-pMHC-I crystal structures were downloaded from the STCRDab[28] database. After removing TCR-pMHC structures in which the peptide contains non-standard amino acids and redundant structures with identical CDR3α, CDR3β, and the presented peptide, we finally obtained 65 non-redundant TCR-pMHC complex structures with a 9-mer peptide bound in the antigen-binding groove of the MHC-I molecule. Then, the contact numbers between CDR3α/β and the bound peptide were calculated using the Python package of PyMOL[40] software. A cutoff distance of 5 Å between each pair of heavy atoms was used for the contact calculations; therefore, two residues are in contact if at least one distance between two heavy atoms is less than the cutoff.

### Potential epitope screening

The mapped epitopes from non-SARS-CoV-2 antigens were utilized to screen potential epitopes against all the possible 9-mer peptides derived from the SARS-CoV-2 proteins (GenBank: MN908947.3). First, the physicochemical similarities in the hotspot region, the fourth to the eighth site, between the mapped epitopes and the SARS-CoV-2 peptides were calculated based on the Euclidean distances of three Atchley factors[41] representing the molecular polarity, size/volume, and electrostatic charge of residues. We chose the Atchley factor, as it was derived from a large number of amino acid indices[41] and has been successfully applied for differentiating disease-associated TCR repertoires[23]. Then, for each mapped epitope, the top 10 most similar SARS-CoV-2 peptides were submitted for immunogenicity prediction. The binding abilities of searched peptides to the top 20 most common HLA class I molecules in China[42] were predicted using NetMHCpan 4.1[43]. In addition, the immunogenicity of searched peptides was predicted and ranked using DeepAntigen[44].

### TCR-pMHC model construction

The structures of TCR-614 and TCR-204 were predicted by a local version of ColabFold[45]. The pMHC models of HLA-B*15:01-NQKLIANQF and HLA-A*3001-KLKTLVATA were constructed by mutating the peptide to the objective epitope based on the crystal structures of pHLA-B*15:01 (PDB id: 6uzq) and pHLA-A*30:01 (PDB id: 6j1w[46]), respectively. Then, the modeled pMHC structures were optimized by 50-ns unbiased MD simulations. Then, the optimized pMHC models were used for molecular docking with TCR to generate TCR-pMHC ternary models. The web server HADDOCK 2.4[47, 48] was utilized to dock TCR to pMHC. The peptide and CDRs were provided as active residues for the docking process. The rigid-body sampling generated 5000 models, and the top 1000 best-scored models were optimized in the semiflexible and water refinement stages. To select the initial model suitable for further analysis, we calculated the contact number between CDR3α and the fourth to the sixth sites in the peptide, as well as the contact number between CDR3β and the fifth to the eighth sites in the peptide, according to the analysis of TCR-pMHC crystal structures. The contact information was calculated with a distance cutoff of 5 Å using the *gmx mindist* command implemented in GROMACS 2020.3 software[49]. Then, a relatively loose threshold of 10 contacts for each CDR3 was used for filtering model candidates. Finally, the best-scored model was selected as the initial structure for the following MD simulations to unveil the TCR-pMHC recognition mechanisms.

### Setups and analysis of MD simulations

The MD simulations were performed using GROMACS-2020 software[49]. The ff14SB[50] force field was used to describe the TCR-pMHC complex, and the TIP3P water model was used to solve the complex. The sodium and chloride ions were added to neutralize the system to an ion concentration of 0.15 M. The cutoff distances of van der Waals (vdW) and short-range electrostatic interactions were set to 12 Å. The Partical-Mesh Ewald[51] (PME) method was used to address the long-range electrostatic interactions. The LINCS[52] algorithm was applied to constrain the chemical bonds. The energy minimization was performed using the steepest descent algorithm, followed by a 200-ps NVT MD simulation with all the protein heavy atoms restrained by a force constant (1000 kJ/mol/nm^2^). The initial velocities of the production MD simulations were randomly assigned at 50 K, and the system was heated to 310 K within 200 ps and kept at 310 K using the velocity rescaling thermostat[53]. Finally, we sampled 100-ns simulation data for the TCR-614-pMHC complex and 200-ns simulation data for the TCR-204-pMHC complex. For each dataset, all the structural analyses were performed based on the last 50-ns simulations.

The *gmx rmsf* command was used to calculate the value of root-mean-square fluctuation. The *gmx select* command was used to calculate the contact number between the TCR and the peptide with a distance cutoff of 6 Å. The HBs were analyzed using the *gmx hbond* command. The solvent-accessible surface area (SASA) was analyzed using FreeSASA[54] software.

## Results

### Investigation of the TCR-pMHC recognition mechanism by leveraging single-cell TCR-seq data and computational tools

Here, we proposed a comprehensive strategy to investigate the molecular mechanisms underlying TCR-pMHC recognition involved in SARS-CoV-2 infection based on available scRNA-seq data (Fig. 1A). First, we collected massive epitope-unknown TCR data, including V(D)J gene usages, CDR3 sequences, and cell clonal expansion information from an scRNA-seq dataset of COVID-19 patients[39]. Then, the T cell clonality analysis was performed to reveal the influences imposed by SARS-CoV-2 infection on T cell immune responses. We also collected epitope-known TCRs with epitope sequences from the VDJdb[27] database. To identify epitopes for epitope-unknown TCRs, we performed sequence-based antigen specificity clustering for epitope-unknown and epitope-known TCRs. For these TCRs that share similar sequence features with epitope-known TCRs targeting SARS-CoV-2 epitopes, we could readily map the corresponding epitopes. For TCRs targeting antigens from other species, we proposed a physicochemical similarity-based strategy to screen potential cross-reactive epitopes against SARS-CoV-2-derived peptides, which allows us to discover neoepitopes. According to the clonality analysis, two representative TCRs with corresponding epitopes were selected for more detailed structural analyses. Finally, we elucidated the epitope recognition mechanisms for both TCRs.

**Figure 1.**
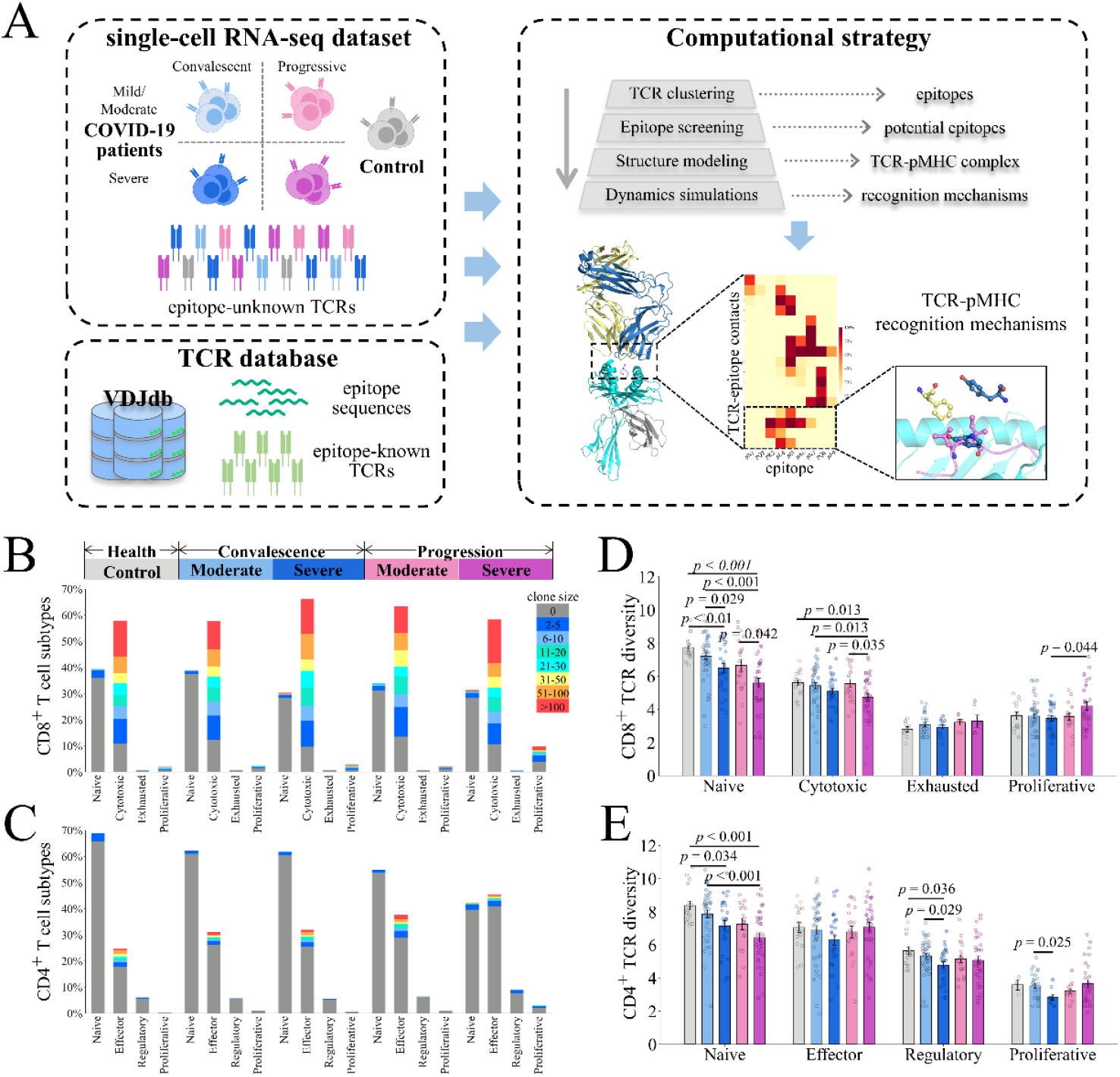
The workflow used to investigate TCR epitope recognition in the current study. (A) A computational strategy to investigate TCR-epitope recognition by leveraging bioinformatics tools. Epitope-unknown TCRs were collected from a massive scRNA-seq dataset. Epitope-known TCRs and epitope sequences were from VDJdb. Representative TCRs were selected based on clone analysis and submitted to the investigation of epitope recognition via tandem computational tools. (B-C) The composition of CD8^+^ T cells (B) and CD4^+^ T cells (C). For each cohort, the histogram indicates the proportion of each subtype and is shown in different colors according to the clone size. (D-E) The TCR diversities of CD8^+^ (D) and CD4^+^ (E) T cell subtypes. Error bars indicate the standard error of the mean, and p-values < 0.05 are labeled above the black line. The p-values were computed using the Mann-Whitney U-test.

### SARS-CoV-2 infection reduces the diversity of the TCR repertoire in cytotoxic CD8^+^ T cells

The epitope-unknown TCR dataset was originally sampled from healthy controls and patients in the disease progression/convalescent stage. Data sampled from patients were also divided by the severity of symptoms into moderate and severe cohorts (Table S2). The collected dataset contains 113,226 (53%) CD4^+^ T cells and 100,529 (47%) CD8^+^ T cells that express αβTCR. According to the original publication, we reannotated the cell subtypes by combining cells similar in the expression of marker genes. We then analyzed the clonal expansion of T cells from healthy controls and patients. We found that CD8^+^ T cells exhibited higher clonal expansion for both controls and patients compared with CD4^+^ T cells (Fig. 1B and 1C). Moreover, for patients with severe symptoms in the disease progression stage, the proportion of proliferative CD8^+^ T cells showed an obvious increment (Fig. 1B), indicating the ongoing expansion of CD8^+^ T cells in these patients. Consistently, a previous study reported that proliferative T cells were elevated significantly in patients and showed associations with COVID-19 severity[39]. For CD4^+^ T cells, naive cells dominated the distribution of T cell subtypes except for severe cohorts in the progression stage (Fig. 1C).

We further delineated the diversity of the TCR repertoire for each cell subtype. Defining TCRs with shared germline gene usage and nucleic acid sequences of CDR3s as identical TCRs, the whole dataset contains ∼8,500 unique TCR sequences that occur more than once. Compared with other cohorts, the diversities of TCR repertoires from severe groups showed significant differences in several subtypes for both CD4^+^ and CD8^+^ T cells (Fig. 1D and 1E). Notably, the diversity of cytotoxic CD8^+^ T cells that consist mainly of clonal T cells decreased significantly for severe cohorts in the progression state (Fig. 1D), indicating the enrichment of antigen-specific TCRs. The shrinkage in the diversity of the TCR repertoire of cytotoxic CD8^+^ T cells probably reflects that the TCR repertoire converges, to some extent, into the SARS-CoV-2-specific spectrum. These results indicated that SARS-CoV-2 infection biased the composition of T cell subtypes and TCR repertoires, especially for patients with severe symptoms in the disease progression stage. Considering the higher clonal expansion and diversity shrinkage in cytotoxic CD8^+^ T cells, we focused on the epitope recognition mechanisms of CD8^+^ T cells in further analysis.

### Identifying SARS-CoV-2-specific TCRs and potential epitopes via a clustering-based pipeline

Antigen specificity is indispensable to comprehending adaptive immune responses, while the lack of epitope information in TCR repertoires hinders the investigation of TCR-epitope recognition. We then proposed a strategy to investigate the antigen specificity and recognition mechanisms for epitope-unknown TCRs. For the epitope-unknown TCR repertoires collected above, we retained only TCR sequences occurring more than once at the amino acid level, resulting in 8,507 epitope-unknown TCRs (Fig. 2A, referred to as dataset UNK in the following sections). In addition, we also collected 1,766 SARS-CoV-2-specific TCRs (Fig. 2A, referred to as dataset VDJ-S) and 29,101 SARS-CoV-2 nonspecific TCRs (referred to as dataset VDJ-N) from VDJdb[27], as well as the corresponding epitope sequences. Then, the epitope information for epitope-unknown TCRs was inferred based on these datasets via two parallel steps. 1) TCR sequence clustering was performed for the UNK dataset and the SARS-CoV-2-specific VDJ-S dataset using GLIPH[20]. For each cluster containing both epitope-known and epitope-unknown TCRs, the epitope sequences were mapped between TCRs (Fig. 2A top). To improve the accuracy of clustering and epitope mapping, we trimmed TCR clusters by retaining only epitope-unknown TCRs directly connected with epitope-known TCRs and vice versa (see details in **Methods**). 2) Identical clustering and additional epitope screening processes were performed for the UNK and SARS-CoV-2 nonspecific datasets VDJ-N (Fig. 2A bottom). As the epitopes in VDJ-N dataset originated from non-SARS-CoV-2 antigens, we then utilized the mapped epitope sequences to screen for similar peptides against SARS-CoV-2 protein sequences. Then, we predicted the binding ability of the searched peptides to 20 common MHC-I molecules using NetMHCpan 4.1[55], followed by immunogenicity prediction and ranking using DeepAntigen[44]. Finally, combining TCR, epitope, and MHC information, we investigated the molecular mechanisms underlying TCR-epitope recognition via structure prediction, molecular docking, and MD simulations (Fig. 2B).

**Figure 2.**
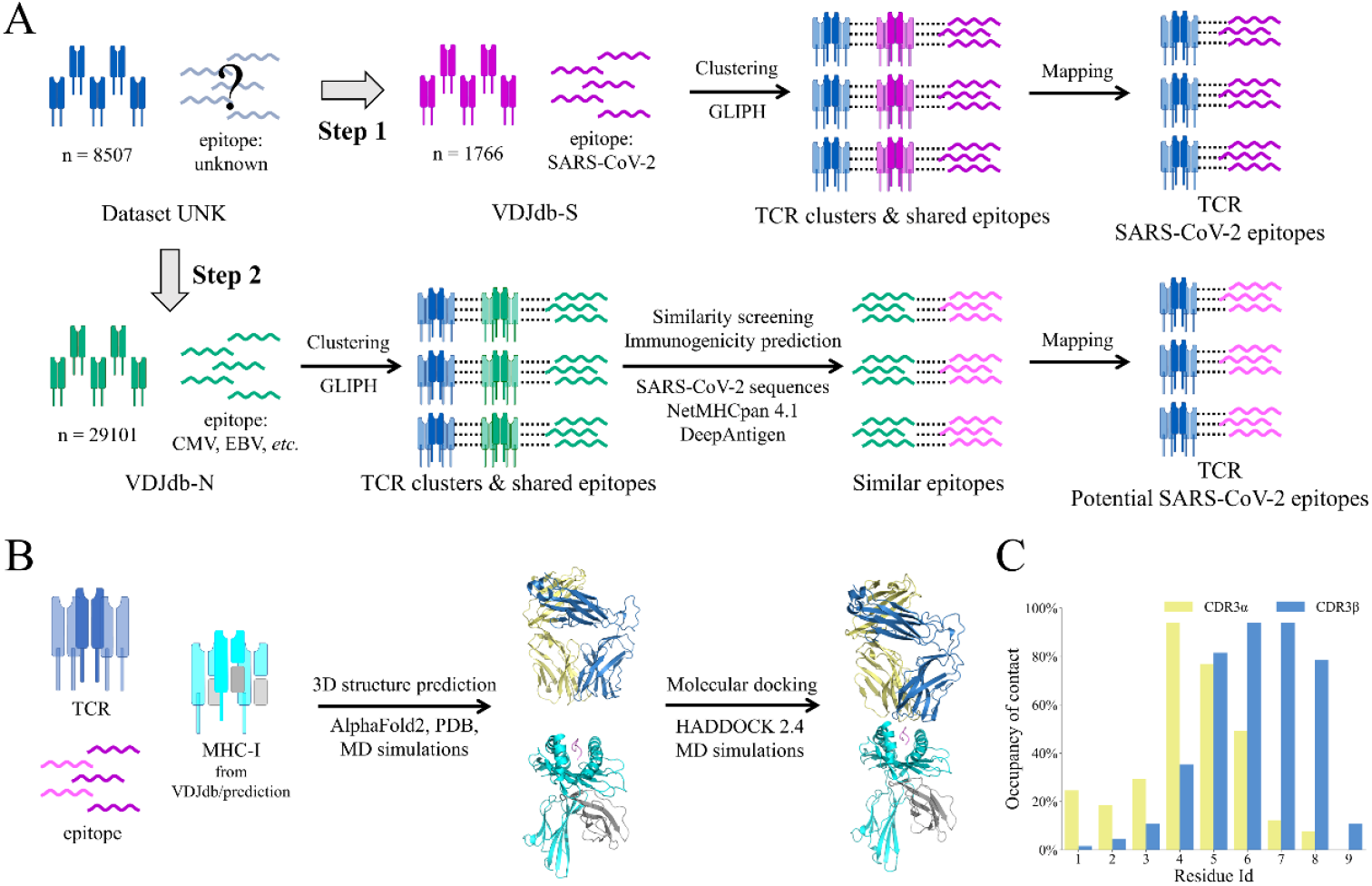
Schematic diagram of the pipeline to identify antigen specificity and explore epitope recognition mechanisms. (A) Flowchart of epitope mapping for epitope-known and epitope-unknown TCRs by sequence clustering. (B) The workflow to investigate the molecular mechanisms of TCR-pMHC recognition. (C) The contact frequency of each site in the 9-mer epitope for 65 TCR-pMHC crystal structures.

Clustering TCRs from the UNK and VDJ-S datasets gave rise to 152 clusters involving 387 epitope-unknown TCRs and 477 epitope-known TCRs targeting SARS-CoV-2 epitopes. Considering the functionality and disease association, we focused on the TCRs carried by highly expanded T cells. Retaining clusters that contain at least one TCR expressed by more than 100 T cells, five clusters were finally identified with epitopes mapped for nineteen epitope-unknown TCRs (Table 1). Four of the five clusters contain TCRs from more than one patient, and none of the clusters contains TCRs from healthy controls. We found that the most-populated TCR (TCR-614, the suffix indicates the number of cells expressing the TCR) clustered in this step was gathered in cluster 1 with a TCR targeting epitope S_919-927_ (NQKLIANQF) from the SARS-CoV-2 spike protein. A sequence motif in CDR3β, ‘SDPE’, was recognized by GLIPH[20] in cluster 1, probably accounting for the shared antigen specificity.

**Table 1.**
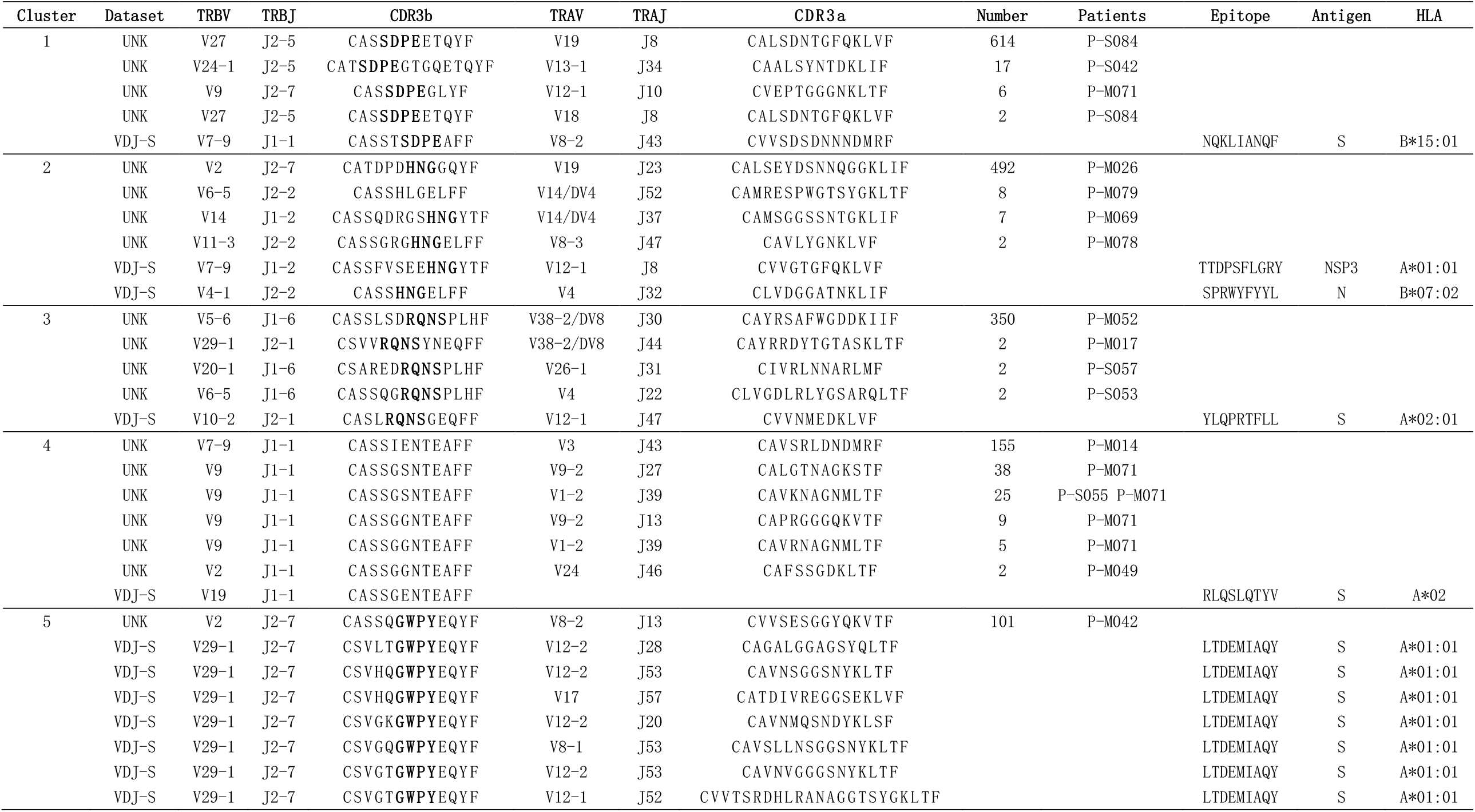
Clusters of epitope-unknown TCRs and SARS-CoV-2-specific TCRs from VDJdb.

In step 2, 2,164 epitope-unknown TCRs and 5,064 TCRs targeting non-SARS-CoV-2 epitopes were gathered into 905 clusters, 11 of which contained at least one TCR expressed by more than 100 T cells (Table 2). A total of eight epitope sequences in the eleven clusters were utilized to screen for potential epitopes against SARS-CoV-2 protein sequences. Prior to the screening, we analyzed the contacts between TCR and the 9-mer antigenic peptide for 65 nonredundant TCR-pMHC complex structures. We found that five hot spots, the fourth to the eighth site, in the presented peptide showed high frequencies in interacting with TCR (Fig. 2C). Therefore, we calculated the physicochemical similarity in the above hot-spot region between the mapped epitopes and all possible 9-mer peptides derived from SARS-CoV-2 proteins to screen for similar peptides. Notably, the most-populated TCR clustered in this step was sampled from healthy controls and shared antigen specificity with a TCR targeting a melanoma-associated neoantigen-derived epitope[56] (Table 2 cluster 1), although it is unclear whether the corresponding donors burden the neoantigen or associated cancers[39]. Thus, we chose the TCR expressed by 204 T cells (TCR-204) in cluster 2 for further analysis. We noticed that six TCRs targeting four distinctive epitopes were highly similar in CDR3β and shared specificity with TCR-204 (Table 2 cluster 2), implying the polyspecificity of TCR-204. Notably, in the potential epitope screening, we found that the shared epitope GLCTLVAML showed high similarity in the TCR-interacting hot spots to a peptide from SARS-CoV-2 NSP3, KLKTLVATA (NSP3_1790-1798_). The peptide was also predicted as a binder to four common HLA alleles (A*02:01, A*02:06, A*30:01, and B*13:02) by NetMHCpan 4.1[43] and ranked with high priority (4/80) in immunogenicity prediction by DeepAntigen[44], suggesting its competence to be an epitope.

**Table 2.**
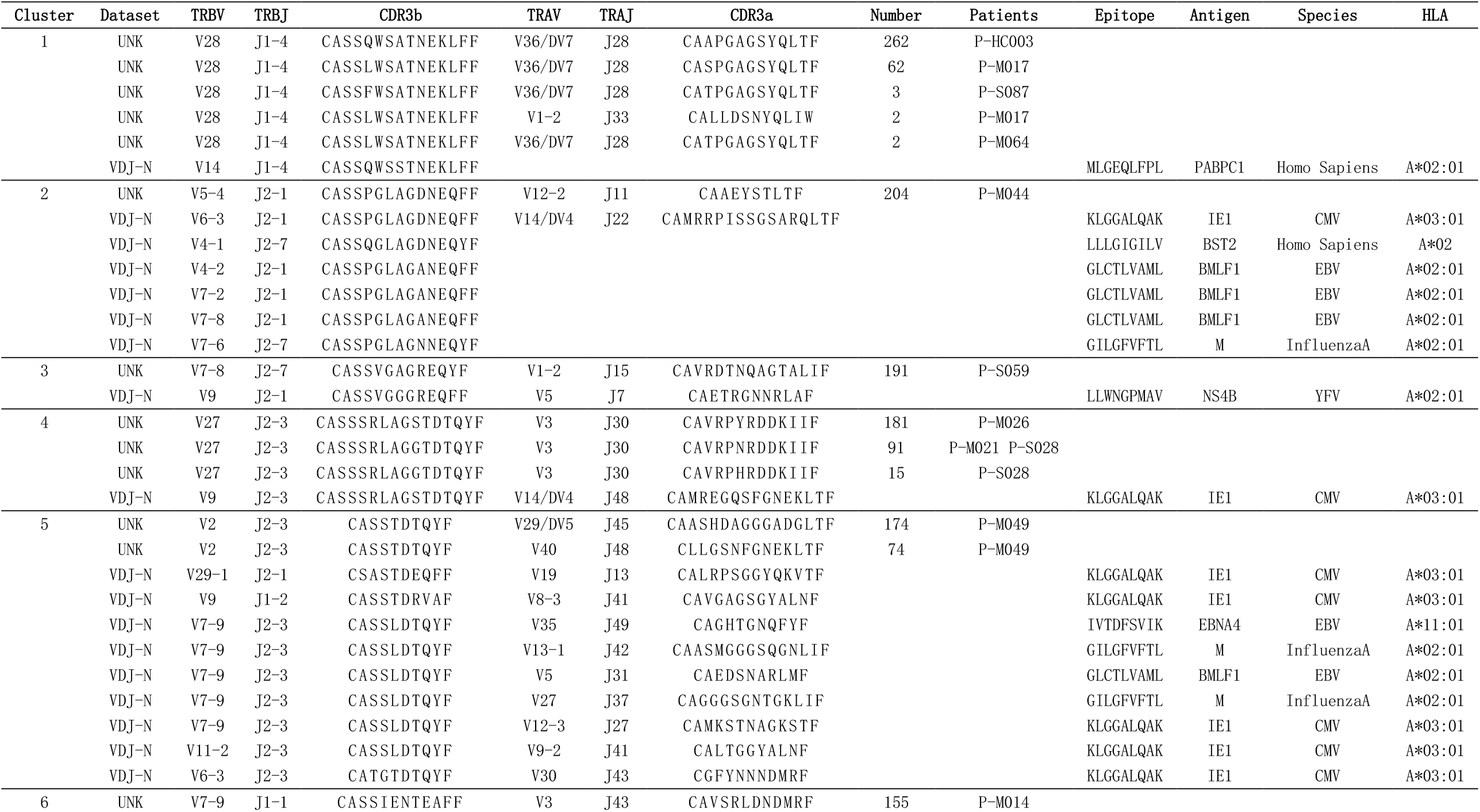
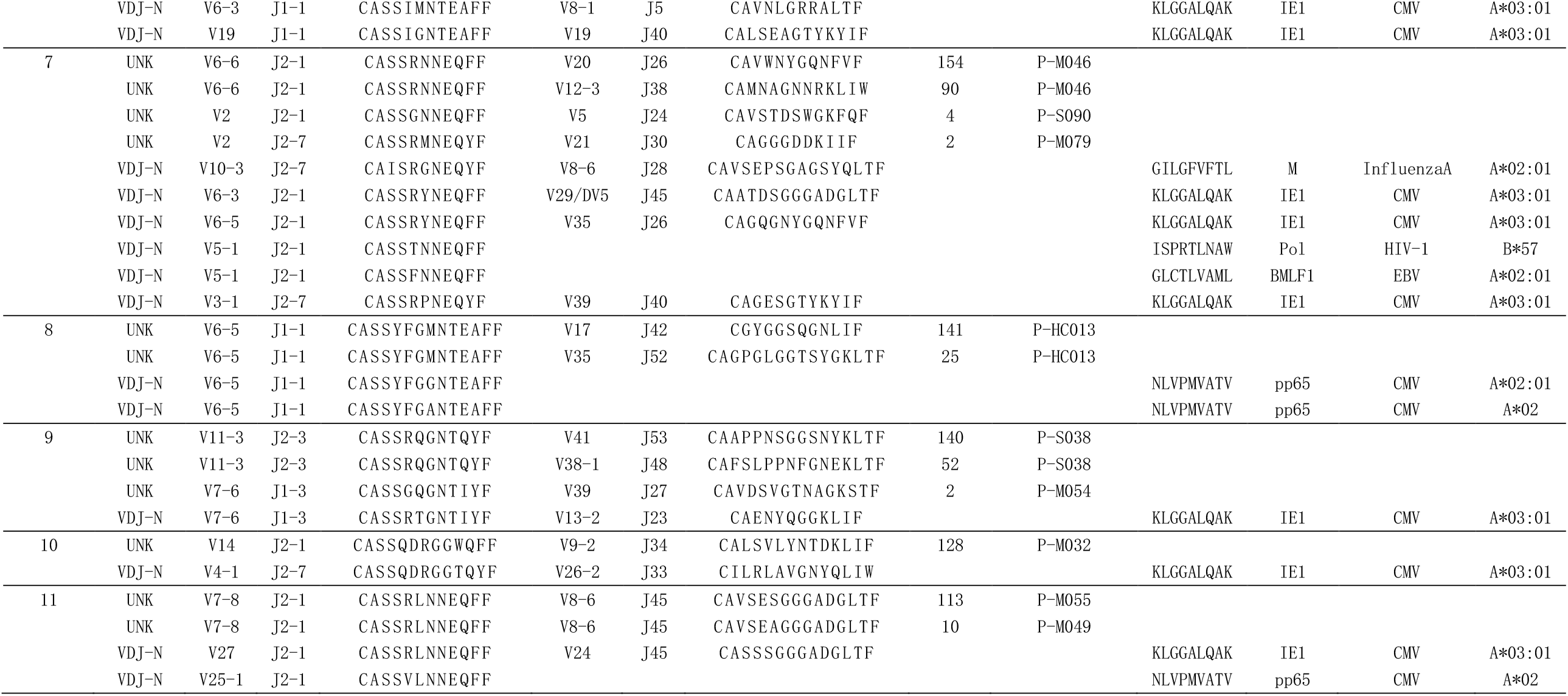
Clusters of epitope-unknown TCRs and non-SARS-CoV-2-specific TCRs from VDJdb.

The identification of reported SARS-CoV-2 epitopes for highly expanded TCRs indicated the capability to identify epitopes for epitope-unknown TCRs via clustering. Meanwhile, combining physicochemical similarity-based epitope screening and immunogenicity prediction, we also discovered neoepitopes for potential treatment. However, the lack of detailed interactions between TCRs and epitopes at the atomic level hinders the understanding and utilization of TCR-pMHC recognition for immunotherapy. Therefore, we further investigated the detailed epitope-recognition mechanisms for the two highly expanded TCRs, TCR-614 and TCR-204.

### TCR-614 employs a hydrophobic clamp and hydrogen bonds to recognize the middle/C-terminus of SARS-CoV-2 epitope S_919-927_

To illustrate the molecular mechanisms of epitope recognition, we constructed the HLA-B*15:01-NQKLIANQF complex model according to the HLA allele information from the VDJdb[27] database and one crystal structure of pHLA-B*15:01 (PDB id: 6uzq). Then, we performed one 50-ns MD simulation to equilibrate the constructed binary complex. The resulting equilibrated model revealed that the presented peptide adopted a canonical conformation in which its two ends were embedded into two pockets in the antigen-binding groove, with pL4-pQ8 exposed to the solvents (Fig. 3A). To stabilize the bound peptide, five and seven hydrogen bonds (HBs) were established between MHC and the peptide N- and C-terminal regions, respectively (Fig. 3B).

**Figure 3.**
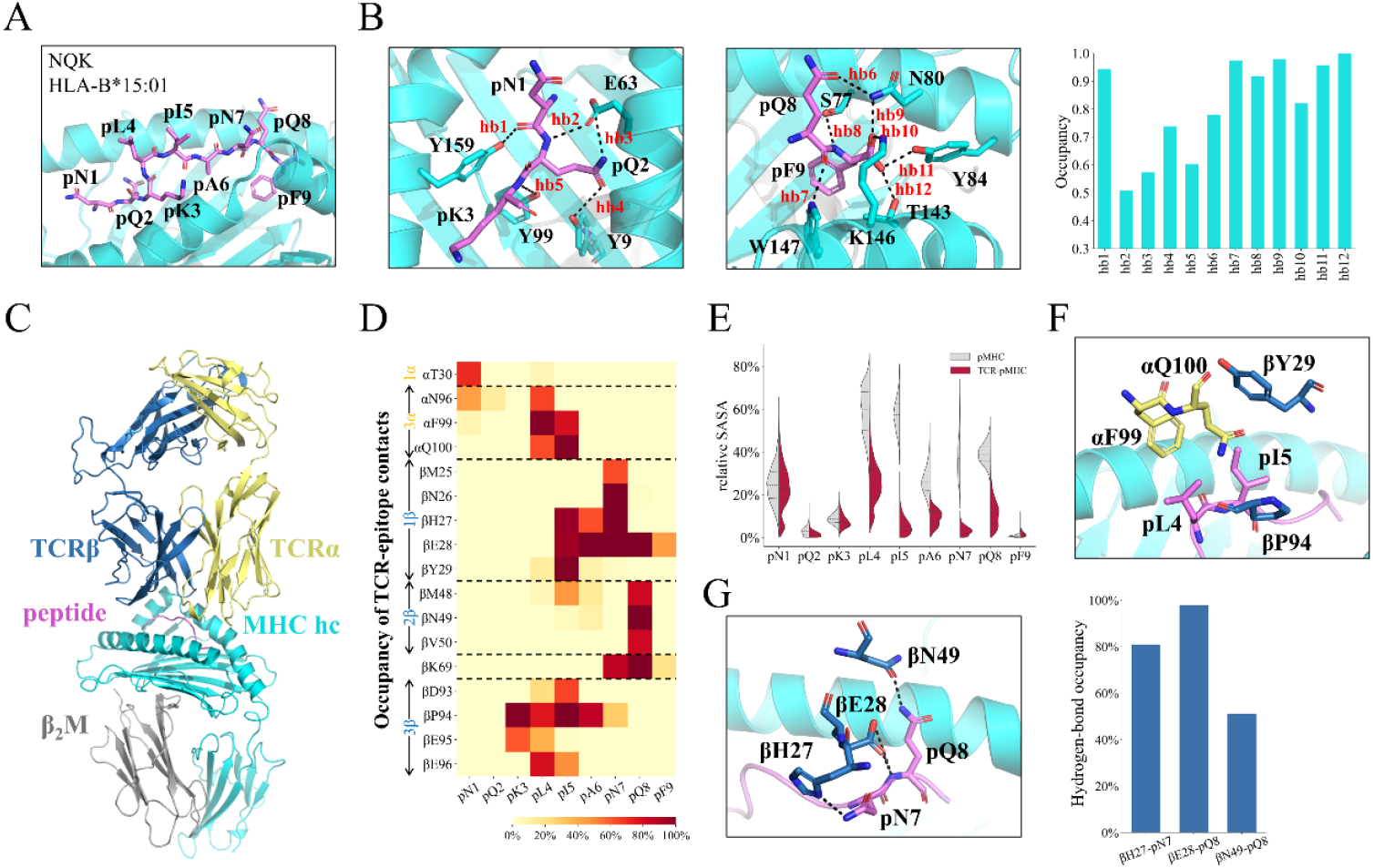
MD simulations reveal the recognition mechanism between the epitope S^919-927^ and TCR-614. (A) The overall conformation of the epitope, NQKLIANQF, presented by HLA-B*15:01. The HLA molecule is shown in the cyan cartoon, and the peptide is highlighted by violet sticks with labels. (B) The HB interactions formed in the peptide N-terminus (left) and C-terminus (middle) and the corresponding occupancies (right). The HBs are indicated by black dashed lines and red labels. (C) The overall structure of the TCR-pMHC complex. (D) The occupancies of contacts formed between the TCR and peptide. The distance cutoff used for the contact calculation is set to 6 Å. (E) Relative solvent-accessible surface area (rSASA) of the peptide during MD simulations for pMHC (gray) and TCR-pMHC (red) complexes. (F) The structural details of the hydrophobic clamp responsible for recognizing pL4pI5. (G) HB interaction networks formed between the TCR β chain and peptide C-terminus (left) and HBs occupancy (right). The HBs are labeled by black dashed lines.

We then sought to construct the ternary complex of the TCR-bound pMHC complex. To this end, we first built the structure of TCR-614 via ColabFold[45] and performed molecular docking to predict the TCR-pMHC model via HADDOCK 2.4[47]. The resulting TCR-pMHC model was then employed as the initial structure for the following 100-ns MD simulations, and the last 50-ns simulation dataset was used for final structural analyses. The equilibrated TCR-pMHC conformation shows that the TCR α and β chain mainly target the peptide N- and C-terminus, respectively (Fig. 3C). In addition, TCR was found to form more direct contacts with the hot spots pL4-pQ8 region (Fig. 3D), leading to a significantly reduced solvent-accessible surface area (SASA) in this region (Fig. 3E). In particular, two peptide residues, pL4 and pI5, could establish more stable interactions with several discontinuous CDR3 residues, including βY29, βD93-E95, and αF99 that clamps the bound peptide, highlighting their importance in TCR recognition (Fig. 3F). Moreover, pN7 and pQ8 could also directly interact with TCR CDR1β and CDR2β, respectively. In addition, several HBs were established between pN7pQ8 and several CDR1/2β residues, i.e., βH27, βE28, and βN49 (Fig. 3G).

### CDR3β conformation is critical for the recognition of the potential SARS-CoV-2 epitope NSP3_1790-1798_ by TCR-204

Likewise, we constructed the TCR-pMHC ternary model for TCR-204 and the potential SARS-CoV-2 epitope KLKTLVATA. To achieve this, we first built the pMHC model based on one crystal structure of pHLA-A*30:01 (PDB id: 6j1w[46]) to which the potential epitope KLKTLVATA was predicted to be a strong binder by NetMHCpan 4.1[43]. Our 50-ns MD simulations indicate that the peptide remained stable and adopted a canonical convex conformation in the antigen-binding groove (Fig. 4A). Similar to the S_919-927_ epitope, the peptide N- and C-terminus were inserted into the antigen-binding groove, with several HBs formed between MHC and the bound peptide (Fig. 4B). In particular, the positively charged pK3 could form salt-bridge interactions with the MHC-E114 (Fig. 4B), which further stabilizes the loaded peptide.

**Figure 4.**
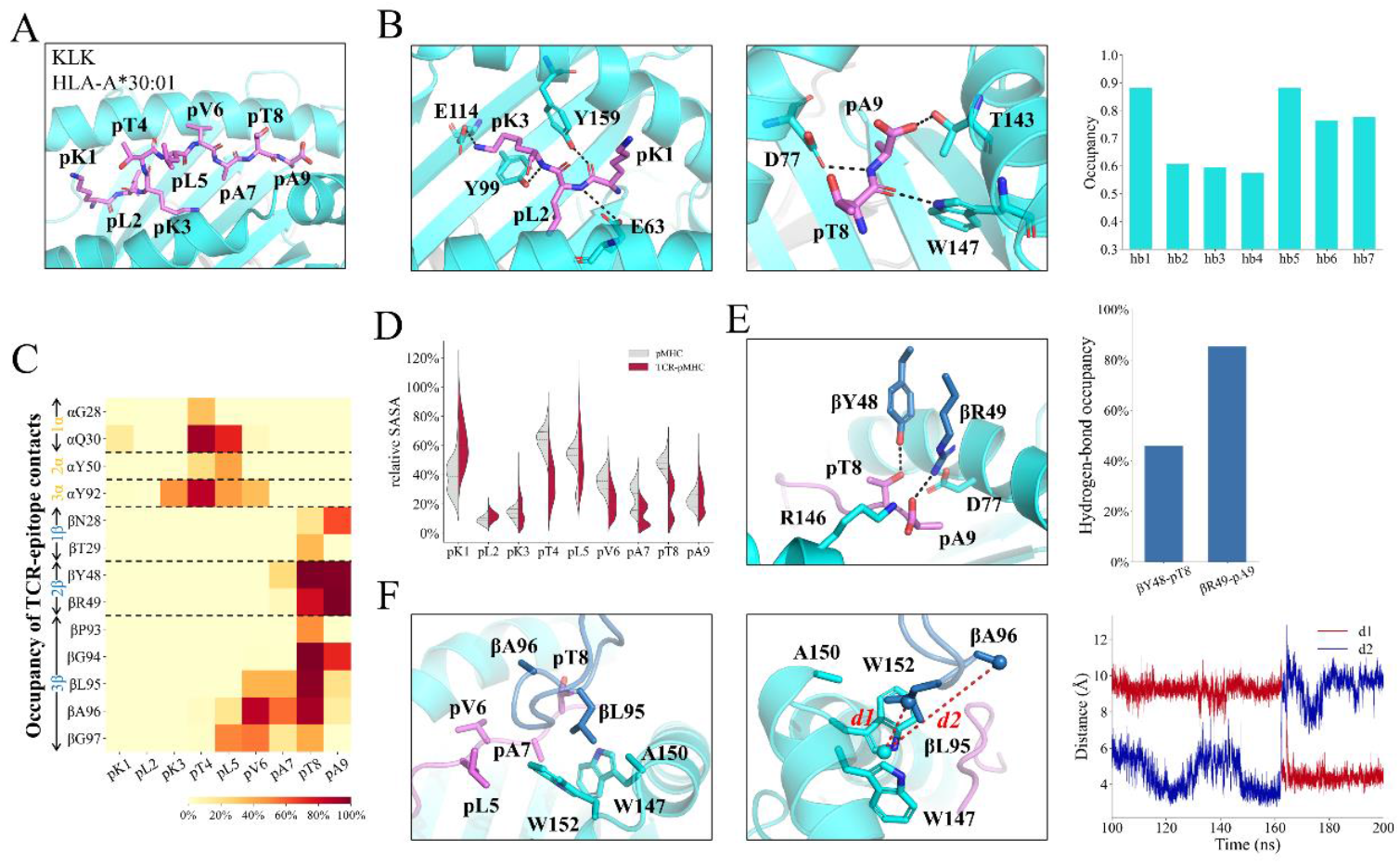
CDR3β of TCR-204 dominates the recognition of epitope NSP3_1790-1798_. (A) The overall conformation of the peptide KLKTLVATA presented by HLA-A*30:01. (B) The hydrogen-bond interactions established between the peptide termini and MHC (left and middle panels) and the corresponding occupancies in MD simulations (right panel). (C) The occupancies of contacts formed between the TCR and peptide. A distance cutoff of 6 Å is used for defining contacts. (D) rSASA of the peptide during MD simulations for pMHC (gray) and TCR-pMHC (red) complexes. (H) HB interactions established between CDR3β and peptide residues (left panel) and the corresponding occupancies (right panel). (F) The hydrophobic core formed by βL95, W147, A150, and W152 (left panel) and two distance measurements indicating the conformational switch of CDR3β (middle and right panels). The distances d1 and d2 are measured between the center of mass (COM) of sidechains of CDR3β residues (βL95 for d1 and βA96 for d2) and MHC residues W147, A150, and W152.

Next, we modeled the structure of TCR-204 via ColabFold[45], and docked this structure to the above pMHC model using HADDOCK 2.4[48]. Then, we performed a 200-ns MD simulation for the ternary TCR-pMHC model to investigate the epitope-recognition mechanisms of NSP3_1790-1798_ by TCR-204. According to the contact calculations between the peptide and TCR, the interacting interface was also located in the hot-spots pT4-pT8 region in which pT8 interacts with a series of CDR3β residues, i.e., βP93-βG97 (Fig. 4C). The SASA analyses also suggest that TCR-204 mainly recognizes the peptide pT4-pT8 region (Fig. 4D). Moreover, the CDR2β residues βY48 and βR49 play critical roles in recognizing the peptide C-terminus via forming HBs, alongside with the MHC residues D77 and R146. In addition to electrostatic interactions, nonpolar interactions are also integral factors in the epitope recognition of NSP3_1790-1798_. Lying over the proximal region of the peptide C-terminus, the hydrophobic region of CDR3β, βP93-G97, simultaneously recognized the peptide and MHC, with the side chain of βL95 embedded into a shallow hydrophobic pocket formed by pA7 and MHC residues W147, A150 & W152 (Fig. 4F *left*). Importantly, the sequence ‘GLAG’ in CDR3β is also highly conserved between TCRs gathered in cluster 2 by GLIPH (Table 2), highlighting its essential role in polyspecificity. Notably, βL95 underwent a conformational shift from a solvent-exposed state to an inserted conformation at ∼165-ns and maintains steady in the remaining simulations (Fig. 4F *middle* and *right*)

## Discussion

In this study, we developed a combinatorial strategy to investigate the disease-associated TCR-pMHC recognition mechanism by leveraging massive single-cell sequencing data and efficient computational tools. The epitope-unknown TCRs from COVID-19 patients were clustered with epitope-known TCRs to identify antigen specificity. According to the single-cell sequencing data, two SARS-CoV-2-associated TCRs, TCR-614 and TCR-204, expressed by highly expanded T cells were subjected to further MD simulations to investigate the molecular mechanisms underlying the TCR and pMHC recognition for two identified epitope sequences NQKLIANQF (S_919-927_) and KLKTLVATA (NSP3_1790-1798_). Combining deep learning-based structure prediction, information-driven docking, and MD simulations, we revealed the critical interactions responsible for epitope recognition by the two TCRs. The CDR3β loops of both TCRs play critical roles in recognizing pMHC molecules, complying well with the conventional understanding of epitope recognition by TCR. For TCR-614, αF99, βY29, and βP94 formed a hydrophobic ‘clamp’ to recognize two hydrophobic peptide residues pL4 and pI5. Several HBs were also formed between CDR1/2β and the peptide C-terminus. For TCR-204, we observed a conformational transition of the CDR3β loop that stabilized the hydrophobic interactions by the insertion of βL95 into a shallow pocket formed by the peptide and MHC residues. Our work provides a computational strategy bridging the single-cell sequencing data of TCRs and structural insights into the epitope-recognition mechanisms. The detailed interactions between the TCR and epitope can be further utilized to facilitate TCR engineering and cancer vaccine design.

The massive TCR sequences generated by high-throughput and single-cell sequencing techniques have greatly promoted the investigation of T cell immune responses. Recently, by clustering experimentally sequenced TCRs with released epitope-known TCRs, researchers were allowed to identify the antigen specificities of epitope-unknown TCRs associated with autoimmune disease[57], viral infection[58], and cancers[59], making it plausible to utilize TCR sequence repertoires for therapeutic discovery. However, the lack of TCR-pMHC complex structures hinders the optimization of TCRs and the rational design of peptide vaccines. To overcome the limitation of structural data, we proposed a computational strategy by combining the TCR clustering method and structural modeling tools, thereby revealing the TCR-pMHC recognition mechanisms for two SARS-CoV-2-associated TCRs at the atomic level. Our MD simulations pinpointed the bilateral residue-residue interactions between TCR and epitope, which is more precise and comprehensive compared with the sole TCR sequence motif recognized by the clustering method. The deep insights into the TCR-pMHC interactions and the accompanying conformational transitions will guide the engineering of TCR-based therapeutics.

Derived from the tumor-specific antigens (TSAs) containing nonsynonymous mutations, the neoepitopes can be discerned and targeted by the immune system, thereby playing a pivotal role in the development of cancer vaccines[60-62]. However, the identification and selection of neoepitopes suitable for vaccine design remain challenging despite several developed strategies[62]. Here, we provided a computational strategy to discover neoepitopes and reveal the structural dynamics underlying epitope recognition by TCRs, which will promote the design and optimization of cancer vaccines.

In this study, we selected two TCRs mainly based on clonal expansion which is a critical process in adaptive immune responses against pathogens[63]. In addition to the extent of clonal expansion, single-cell approaches can simultaneously capture TCR sequences coupled with more features, such as gene/protein expression and chromatin accessibility[26]. These features can also be utilized to identify the TCRs carried by functional or characteristic T cells for investigating TCR-pMHC recognition by structural modeling. Based on the constructed TCR-pMHC model, researchers could further improve the binding affinity and specificity of TCRs by combining other computational methods[30, 31, 33]. For the epitope, the recognized peptide repertoire by a certain TCR is highly expanded owing to the polyspecificity. Here, we determined a hot-spots region, position 4 to position 8 of the 9-mer peptide, which was subsequently used for potential epitope screening by physicochemical similarity searching against SARS-CoV-2-derived peptides. Consistent with previous studies, the cross-reactive peptides exhibited similarity in hot-spots regions[64, 65], demonstrating the feasibility of discovering cross-reactive peptides by similarity searching. However, even highly distinctive peptides can be recognized by the same TCR with different conformations[64], indicating the limited efficacy of similarity searching in the discovery of cross-reactive peptides. In addition, a variety of post-translational modifications can reshape the peptides presented by MHC molecules and influence the T cell immune response[66], which further complicates TCR-pMHC recognition. To better understand and utilize T cell immune responses, the molecular basis underlying TCR-pMHC recognition remains to be comprehensively elucidated in the future, and further studies are necessary to unveil the association between cellular characteristics and TCR-pMHC interactions.

## Data Availability

The epitope-unknown TCR data were collected from a massive scRNA-seq dataset[39] and can be downloaded from the NCBI GEO database (GES158055). The epitope-known TCRs can be downloaded from the VDJdb[27] database. The TCR-pMHC complex structures can be downloaded from the STCRDab[28] database. The datasets used and/or analysed during the current study are available from the corresponding author on reasonable request.

## Authors’ contributions

XZ, L-TD, JH, and KS designed the project. KS and YS performed bioinformatic analyses and simulations. HX and YS provided important assistance for the interpretation of the data. KS, XZ, and L-TD wrote the manuscript. All authors read and approved the final manuscript.

## Acknowledgements

We gratefully acknowledge the computational support from High-Performance Computing of Shanghai Jiao Tong University.

## References

1. Fooksman DR, Vardhana S, Vasiliver-Shamis G et al. Functional anatomy of T cell activation and synapse formation, Annu Rev Immunol 2010;28:79–105.

2. Kumar BV, Connors TJ, Farber DL. Human T Cell Development, Localization, and Function throughout Life, Immunity 2018;48:202–213.

3. Liu B, Kolawole EM, Evavold BD. Mechanobiology of T Cell Activation: To Catch a Bond, Annu Rev Cell Dev Biol 2021;37:65–87.

4. Bassing CH, Swat W, Alt FW. The mechanism and regulation of chromosomal V(D)J recombination, Cell 2002;109 Suppl:S45–55.

5. Krangel MS. Mechanics of T cell receptor gene rearrangement, Curr Opin Immunol 2009;21:133–139.

6. Dupic T, Marcou Q, Walczak AM et al. Genesis of the αβ T-cell receptor, PLoS Comput Biol 2019;15:e1006874.

7. Davis MM, Boyd SD. Recent progress in the analysis of αβT cell and B cell receptor repertoires, Curr Opin Immunol 2019;59:109–114.

8. Robins HS, Campregher PV, Srivastava SK et al. Comprehensive assessment of T-cell receptor beta-chain diversity in alphabeta T cells, Blood 2009;114:4099–4107.

9. Wucherpfennig KW, Allen PM, Celada F et al. Polyspecificity of T cell and B cell receptor recognition, Semin Immunol 2007;19:216–224.

10. Sewell AK. Why must T cells be cross-reactive?, Nat Rev Immunol 2012;12:669–677.

11. Lowe KL, Cole D, Kenefeck R et al. Novel TCR-based biologics: mobilising T cells to warm ‘cold’ tumours, Cancer Treat Rev 2019;77:35–43.

12. Robins HS, Srivastava SK, Campregher PV et al. Overlap and effective size of the human CD8+ T cell receptor repertoire, Sci Transl Med 2010;2:47ra64.

13. Chiffelle J, Genolet R, Perez MA et al. T-cell repertoire analysis and metrics of diversity and clonality, Curr Opin Biotechnol 2020;65:284–295.

14. Bradley P, Thomas PG. Using T Cell Receptor Repertoires to Understand the Principles of Adaptive Immune Recognition, Annu Rev Immunol 2019;37:547–570.

15. Krummey SM, Morris AB, Jacobs JR et al. CD45RB Status of CD8(+) T Cell Memory Defines T Cell Receptor Affinity and Persistence, Cell Rep 2020;30:1282-1291.e1285.

16. Schober K, Voit F, Grassmann S et al. Reverse TCR repertoire evolution toward dominant low-affinity clones during chronic CMV infection, Nat Immunol 2020;21:434–441.

17. Pruessmann W, Rytlewski J, Wilmott J et al. Molecular analysis of primary melanoma T cells identifies patients at risk for metastatic recurrence, Nat Cancer 2020;1:197–209.

18. Valpione S, Galvani E, Tweedy J et al. Immune-awakening revealed by peripheral T cell dynamics after one cycle of immunotherapy, Nat Cancer 2020;1:210–221.

19. Dash P, Fiore-Gartland AJ, Hertz T et al. Quantifiable predictive features define epitope-specific T cell receptor repertoires, Nature 2017;547:89–93.

20. Glanville J, Huang H, Nau A et al. Identifying specificity groups in the T cell receptor repertoire, Nature 2017;547:94–98.

21. Pogorelyy MV, Minervina AA, Shugay M et al. Detecting T cell receptors involved in immune responses from single repertoire snapshots, PLoS Biol 2019;17:e3000314.

22. Zhang H, Liu L, Zhang J et al. Investigation of Antigen-Specific T-Cell Receptor Clusters in Human Cancers, Clin Cancer Res 2020;26:1359–1371.

23. Ostmeyer J, Christley S, Toby IT et al. Biophysicochemical Motifs in T-cell Receptor Sequences Distinguish Repertoires from Tumor-Infiltrating Lymphocyte and Adjacent Healthy Tissue, Cancer Res 2019;79:1671–1680.

24. Huang H, Wang C, Rubelt F et al. Analyzing the Mycobacterium tuberculosis immune response by T-cell receptor clustering with GLIPH2 and genome-wide antigen screening, Nat Biotechnol 2020;38:1194–1202.

25. Zhang H, Zhan X, Li B. GIANA allows computationally-efficient TCR clustering and multi-disease repertoire classification by isometric transformation, Nat Commun 2021;12:4699.

26. Pai JA, Satpathy AT. High-throughput and single-cell T cell receptor sequencing technologies, Nat Methods 2021;18:881–892.

27. Shugay M, Bagaev DV, Zvyagin IV et al. VDJdb: a curated database of T-cell receptor sequences with known antigen specificity, Nucleic Acids Res 2018;46:D419–d427.

28. Leem J, de Oliveira SHP, Krawczyk K et al. STCRDab: the structural T-cell receptor database, Nucleic Acids Res 2018;46:D406–d412.

29. Rossjohn J, Gras S, Miles JJ et al. T cell antigen receptor recognition of antigen-presenting molecules, Annu Rev Immunol 2015;33:169–200.

30. Pierce BG, Hellman LM, Hossain M et al. Computational design of the affinity and specificity of a therapeutic T cell receptor, PLoS Comput Biol 2014;10:e1003478.

31. Hellman LM, Foley KC, Singh NK et al. Improving T Cell Receptor On-Target Specificity via Structure-Guided Design, Mol Ther 2019;27:300–313.

32. Rosenberg AM, Baker BM. Engineering the T cell receptor for fun and profit: Uncovering complex biology, interrogating the immune system, and targeting disease, Curr Opin Struct Biol 2022;74:102358.

33. Crean RM, Pudney CR, Cole DK et al. Reliable In Silico Ranking of Engineered Therapeutic TCR Binding Affinities with MMPB/GBSA, J Chem Inf Model 2022;62:577–590.

34. Baek M, DiMaio F, Anishchenko I et al. Accurate prediction of protein structures and interactions using a three-track neural network, Science 2021;373:871–876.

35. Jumper J, Evans R, Pritzel A et al. Highly accurate protein structure prediction with AlphaFold, Nature 2021;596:583–589.

36. Ambrosetti F, Jiménez-GarcÁa B, Roel-Touris J et al. Modeling Antibody-Antigen Complexes by Information-Driven Docking, Structure 2020;28:119-129.e112.

37. Peacock T, Chain B. Information-Driven Docking for TCR-pMHC Complex Prediction, Front Immunol 2021;12:686127.

38. Wu P, Zhang T, Liu B et al. Mechano-regulation of Peptide-MHC Class I Conformations Determines TCR Antigen Recognition, Mol Cell 2019;73:1015-1027.e1017.

39. Ren X, Wen W, Fan X et al. COVID-19 immune features revealed by a large-scale single-cell transcriptome atlas, Cell 2021;184:1895-1913.e1819.

40. Schrodinger, LLC. The PyMOL Molecular Graphics System, Version 1.8. 2015.

41. Atchley WR, Zhao J, Fernandes AD et al. Solving the protein sequence metric problem, Proc Natl Acad Sci U S A 2005;102:6395–6400.

42. He Y, Li J, Mao W et al. HLA common and well-documented alleles in China, Hla 2018;92:199–205.

43. Reynisson B, Alvarez B, Paul S et al. NetMHCpan-4.1 and NetMHCIIpan-4.0: improved predictions of MHC antigen presentation by concurrent motif deconvolution and integration of MS MHC eluted ligand data, Nucleic Acids Res 2020;48:W449–w454.

44. Shi Y, Guo Z, Su X et al. DeepAntigen: a novel method for neoantigen prioritization via 3D genome and deep sparse learning, Bioinformatics 2020;36:4894–4901.

45. Mirdita M, Schütze K, Moriwaki Y et al. ColabFold: making protein folding accessible to all, Nat Methods 2022;19:679–682.

46. Zhu S, Liu K, Chai Y et al. Divergent Peptide Presentations of HLA-A(*)30 Alleles Revealed by Structures With Pathogen Peptides, Front Immunol 2019;10:1709.

47. Dominguez C, Boelens R, Bonvin AM. HADDOCK: a protein-protein docking approach based on biochemical or biophysical information, J Am Chem Soc 2003;125:1731–1737.

48. van Zundert Gcp, Rodrigues J, Trellet M et al. The HADDOCK2.2 Web Server: User-Friendly Integrative Modeling of Biomolecular Complexes, J Mol Biol 2016;428:720–725.

49. Van Der Spoel D, Lindahl E, Hess B et al. GROMACS: fast, flexible, and free, J Comput Chem 2005;26:1701–1718.

50. Maier JA, Martinez C, Kasavajhala K et al. ff14SB: Improving the Accuracy of Protein Side Chain and Backbone Parameters from ff99SB, J Chem Theory Comput 2015;11:3696–3713.

51. Essmann U, Perera L, Berkowitz ML et al. A smooth particle mesh Ewald method, Journal of Chemical Physics 1995;103:8577–8593.

52. Hess B, Bekker H, Berendsen HJC et al. LINCS: A linear constraint solver for molecular simulations, Journal of Computational Chemistry 1997;18:1463–1472.

53. Bussi G, Donadio D, Parrinello M. Canonical sampling through velocity rescaling, Journal of Chemical Physics 2007;126.

54. Mitternacht S. FreeSASA: An open source C library for solvent accessible surface area calculations, F1000Res 2016;5:189.

55. Jensen KK, Andreatta M, Marcatili P et al. Improved methods for predicting peptide binding affinity to MHC class II molecules, Immunology 2018;154:394–406.

56. Carreno BM, Magrini V, Becker-Hapak M et al. Cancer immunotherapy. A dendritic cell vaccine increases the breadth and diversity of melanoma neoantigen-specific T cells, Science 2015;348:803–808.

57. Akama-Garren EH, van den Broek T, Simoni L et al. Follicular T cells are clonally and transcriptionally distinct in B cell-driven mouse autoimmune disease, Nat Commun 2021;12:6687.

58. Schneider-Hohendorf T, Gerdes LA, Pignolet B et al. Broader Epstein-Barr virus-specific T cell receptor repertoire in patients with multiple sclerosis, J Exp Med 2022;219.

59. Goncharov MM, Bryushkova EA, Sharaev NI et al. Pinpointing the tumor-specific T cells via TCR clusters, Elife 2022;11.

60. Shemesh CS, Hsu JC, Hosseini I et al. Personalized Cancer Vaccines: Clinical Landscape, Challenges, and Opportunities, Mol Ther 2021;29:555–570.

61. Lin MJ, Svensson-Arvelund J, Lubitz GS et al. Cancer vaccines: the next immunotherapy frontier, Nat Cancer 2022;3:911–926.

62. Sellars MC, Wu CJ, Fritsch EF. Cancer vaccines: Building a bridge over troubled waters, Cell 2022;185:2770–2788.

63. Adams NM, Grassmann S, Sun JC. Clonal expansion of innate and adaptive lymphocytes, Nat Rev Immunol 2020;20:694–707.

64. Riley TP, Hellman LM, Gee MH et al. T cell receptor cross-reactivity expanded by dramatic peptide-MHC adaptability, Nat Chem Biol 2018;14:934–942.

65. Adams JJ, Narayanan S, Birnbaum ME et al. Structural interplay between germline interactions and adaptive recognition determines the bandwidth of TCR-peptide-MHC cross-reactivity, Nat Immunol 2016;17:87–94.

66. Kacen A, Javitt A, Kramer MP et al. Post-translational modifications reshape the antigenic landscape of the MHC I immunopeptidome in tumors, Nat Biotechnol 2022.

